# Acoustically patterned hepatic lobule-like units for vascularized artificial liver

**DOI:** 10.64898/2026.09.21.753064

**Authors:** Huong Thi Le, Van Phu Le, Hien Thi Le, Chanh Trung Nguyen, Andreas Lenshof, Chaenyung Cha, Sung Hoon Back, Thomas Laurell, Kyo-in Koo

## Abstract

Engineering a transplantable artificial liver requires reconstituting the hepatic lobule, whose densely cellular parenchyma is organized around a hierarchically branched vascular network. Achieving high cellular density and hierarchical vascularization within a single fabrication step remains a major challenge. Here, we present an ultrasonic standing-wave (USW)-based strategy that concurrently organizes vascular and parenchymal compartments during a single extrusion. HepG2–endothelial cell aggregates were fabricated as building blocks and co-extruded with single endothelial cells through a glass capillary under USW actuation in a liver-derived decellularized extracellular matrix–alginate hydrogel, enabling rapid, ultraviolet-free crosslinking. Due to their size difference, the single cells were patterned from one supplying line in the acoustic pressure node to four microvascular lines while the aggregates localized away from the acoustic pressure node, reconstituting the basic microvascular–parenchymal unit of the lobule. Aggregate size increased with seeding density while maintaining uniformity and over 90% viability, and the patterned endothelial cells formed perfusable lumina through which 5 µm microparticles flowed freely. Compared with hepatocyte-only cultures, the vascularized constructs showed a 1.7-fold increase in urea production, and 2.7- and 6.2-fold increases in *CYP1A2* and *CYP3A4* expression, respectively. Following implantation into the mouse liver, the scaffolds showed favorable biocompatibility, sustained proliferation over 21 days, and progressive host integration, with the smallest blood-containing lumina maturing from 26.1 µm to 7.9 µm in diameter and increasing collagen deposition at the graft–host interface. This approach provides a template-free route to hierarchically vascularized hepatic units and a promising step toward implantable liver tissue.

## 1. Introduction

The liver performs a wide range of vital functions, including protein synthesis, urea metabolism, and xenobiotic detoxification (Huppert & Schwartz, Annu Rev Physiol, 2023). End-stage liver disease is a major cause of mortality worldwide, and orthotopic liver transplantation remains the only definitive treatment. However, the demand for donor organs far exceeds the available supply, leaving many patients on waiting lists (Mazza et al., Hepatol Commun, 2018; Terrault et al., Clinical Gastroenterology and Hepatology, 2023). This persistent shortage has motivated intensive efforts in liver tissue engineering to construct transplantable, functional hepatic tissues *in vitro* (Shi et al., Biomolecules, 2024). Among the various biomaterials explored for this purpose, liver-derived decellularized extracellular matrix (dECM) has emerged as a particularly attractive substrate, as it preserves tissue-specific matrix components and has been shown to enhance hepatocyte function and support hepatic differentiation (Lee et al., Biomacromolecules, 2017; Uygun et al., Nature Medicine, 2010).

A central challenge in engineering functional liver tissue is to reconstitute the hepatic lobule, the basic structural and functional unit of the liver. Each lobule comprises a densely packed hepatocyte compartment organized around a hierarchical vascular network, in which blood flows from the periportal microvessels through the sinusoids toward the central vein (Kietzmann, Redox Biol, 2017; Trefts et al., Curr Biol, 2017). Reproducing this organization *in vitro* requires both a high cellular density (over 1 × 10^8^ cells/mL (Daly et al., Cell, 2021)) and tissue-level cellular organization comparable to those of the native organ. Conventional bioinks, in which cells are dispersed as individual cells within a hydrogel, are limited in the cell density (under 1 × 10^7^ cells/mL (Daly et al., Cell, 2021)) they can accommodate while maintaining printability, which constrains the formation of functional tissue (Duong et al., Eur Cell Mater, 2019; Fang et al., Biofabrication, 2023; Jeon et al., Gut and Liver, 2017). To address this limitation, the assembly of pre-formed multicellular building blocks such as spheroids, organoids, and cell aggregates, collectively referred to as organ building blocks (OBBs) has emerged as a powerful strategy. Because each building block already contains thousands of cells organized at the tissue scale, OBB-based approaches can rapidly generate constructs with the requisite cellular density, microarchitecture, and function (Skylar-Scott et al., Science Advances, 2019). In the context of the liver, the aggregation of hepatocytes enhances cell-cell interactions and promotes the maintenance of hepatic phenotype and function relative to dispersed-cell cultures (Fang et al., Biofabrication, 2023), making hepatocyte-based aggregates promising modular units for reconstructing the hepatocyte compartment of the lobule.

Beyond cellular density, the generation of implantable-scale tissues critically depends on the incorporation of a well-developed vascular network. In the absence of perfusable vasculature, oxygen and nutrient diffusion restricts the viability of engineered tissues to a thickness of only approximately 100 - 200 µm, beyond which the tissue core becomes necrotic (Duong et al., Biofabrication, 2020; Grebenyuk & Ranga, Frontiers in Bioengineering and Biotechnology, 2019; Nguyen et al., Int J Bioprint, 2022). The hepatic lobule exemplifies the hierarchical vascular organization required to overcome this limitation: a continuous network extending from larger supplying and draining vessels, through branched sinusoidal microvessels, down to the immediate perivascular space surrounding each hepatocyte (Aird, Circulation Research, 2007; Kietzmann, Redox Biol, 2017). To sustain tissues of clinically relevant size, an engineered vasculature must similarly be hierarchically organized, such that larger supplying vessels capable of integrating with the host circulation connect to branched microvascular networks, which in turn perfuse the interior of the densely cellular building blocks (Landau et al., Development, 2024). Importantly, engineered endothelial networks have been shown to anastomose with the host vasculature after implantation, enabling rapid perfusion of the graft (Brady et al., Scientific Reports, 2023; Duong et al., Biomaterials, 2025).

Diverse biofabrication strategies have sought to recapitulate the hepatic lobule, including cell spheroids, organoids, microfluidic devices, and bioprinting-based approaches (Bhise et al., Biofabrication, 2016; Coll et al., Cell Stem Cell, 2018; Wu et al., Journal of Hepatology, 2019). While these approaches have reproduced aspects of lobular organization, vascularization has typically relied on sacrificial templating, coaxial extrusion, laser patterning, or microfluidic self-assembly of endothelial networks. These methods often require sacrificial templates, specialized printing hardware, or ultraviolet exposure, and provide limited control over the hierarchical branching that characterizes the lobular microvasculature. Moreover, most OBB-based fabrication methods focus on achieving high cellular density but do not simultaneously generate an integrated, hierarchically branched vascular structure within the same process. As a result, the concurrent formation of a densely cellular hepatic compartment and a hierarchically organized vascular compartment within a single fabrication step remains a significant challenge.

To address this challenge, we developed an ultrasound standing-wave (USW)-based strategy that integrates hepatocyte-endothelial cell building blocks with acoustically patterned vascular structures within a single extrusion process. We first fabricated HepG2-endothelial cell (HG-EC) aggregates as functional building blocks, and then co-extruded these aggregates with single endothelial cells through a glass capillary under USW actuation. In this process, liver-derived dECM served as the primary matrix to support hepatic function, while alginate was incorporated to enable rapid, UV-free crosslinking of the construct. Owing to the size difference single cells and aggregates, the single endothelial cells were focused at the pressure nodes to form branched vascular patterns, ranging from a single supplying line to four branched microvascular lines, while the aggregates localized to the surrounding regions (Augustsson et al., Anal Chem, 2012; Petersson et al., Analyst, 2004). This configuration reconstitutes the basic microvascular-parenchymal unit of the hepatic lobule (branched microvessels surrounded by hepatocyte clusters). This approach builds upon our previous demonstrations of USW-mediated patterning of endothelial cells (Le et al., Biofabrication, 2023) and fibroblasts (Koo et al., Micromachines, 2021) into network-like structures, and extends the concept to a functional, vascularized liver tissue unit. We further evaluated the vascularization, hepatic function, and *in vivo* integration of the resulting scaffolds following implantation into the mouse liver.

## 2. Materials and methods

### 2.1. Theory and concept

A 2 MHz USW in two types of square glass capillaries (400 µm and 800 µm) enables a transition between one focused cell stream and four cell streams of single cells. When applying 2 MHz ultrasound in a 400 µm square-shaped glass capillary, a half-wavelength standing wave was established, generating a single pressure node at the center of the channel. When the capillary dimension was doubled to 800 µm, the square cross-section supported full-wavelength resonance along both the *x*- and *y*-directions, producing four pressure nodes located λ/4 from the channel side walls.

When single cells were mixed with cell aggregates, the two different-sized objects were driven in opposite directions owing to differences in their acoustic contrast factor (□). The single cells, which exhibit a positive contrast factor, were driven toward the pressure nodes, whereas the cell aggregates were driven toward the pressure antinodes. This occurs when the size of the cell aggregate is a significant fraction of the wavelength and thus the fundamental criterion for the acoustic radiation force within the Rayleigh limit (particle radius << wavelength /2π) no longer applies (Silva et al., Physical Review Applied, 2019) (Fig. 1 and Supplementary video 1).

**Figure 1.**
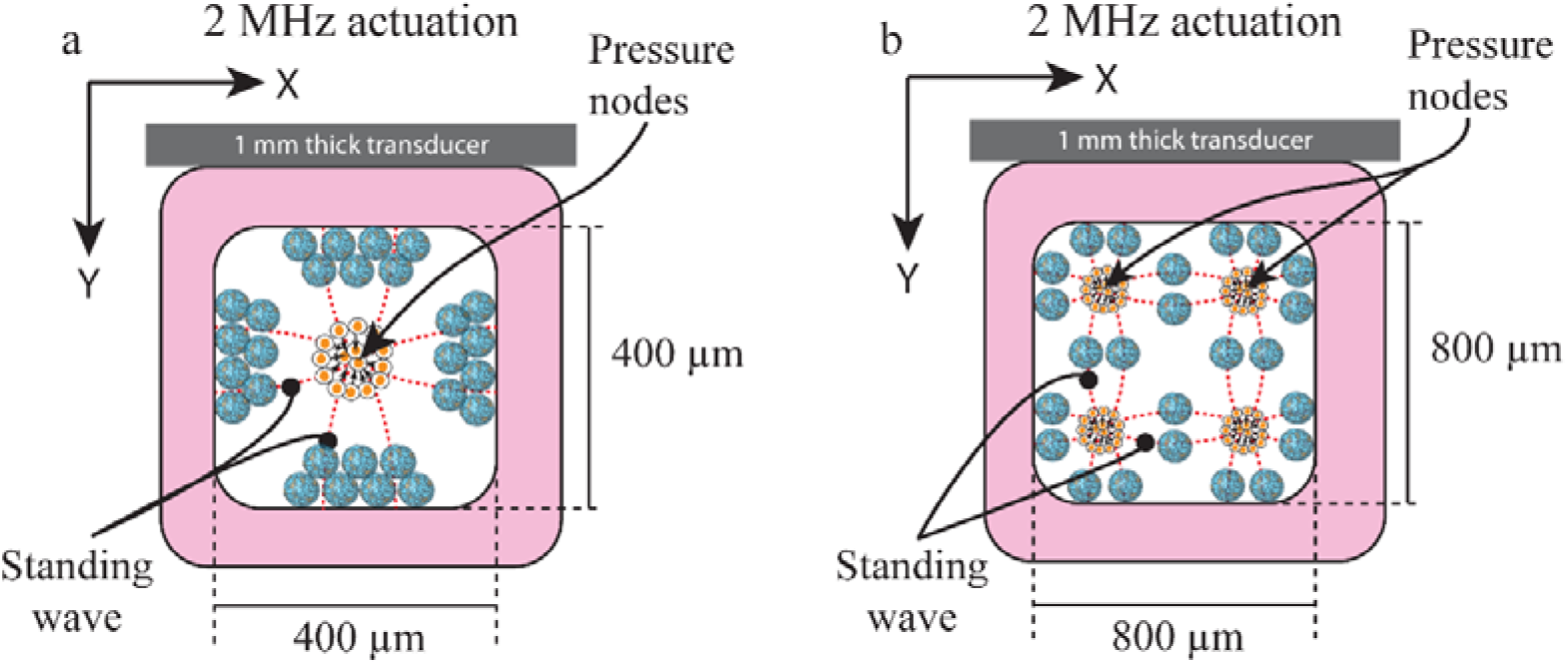
Schematic illustration of the standing-wave patterns in the capillary cross-section upon actuation (a) at 2 MHz in a 400 μm square glass capillary, yielding a single pressure node at the center toward individual cells move, and (b) at 2 MHz in the 800 μm square-shaped glass capillary, producing four pressure nodes, one in each quadrant of the cross section, toward which individual cells move. The cell aggregates, in contrast, are driven toward the pressure antinodes. The transducers and glass capillaries are not drawn to scale.

This spatial separation enables endothelial cells to be patterned into a branched configuration, with HepG2-endothelial cell aggregates surrounded these branched endothelial cells (Fig. 2). The fabricated scaffolds were subsequently cultured in the presence of alginate lyase to promote microvascular network formation and to enhance hepatocyte function.

**Figure 2.**
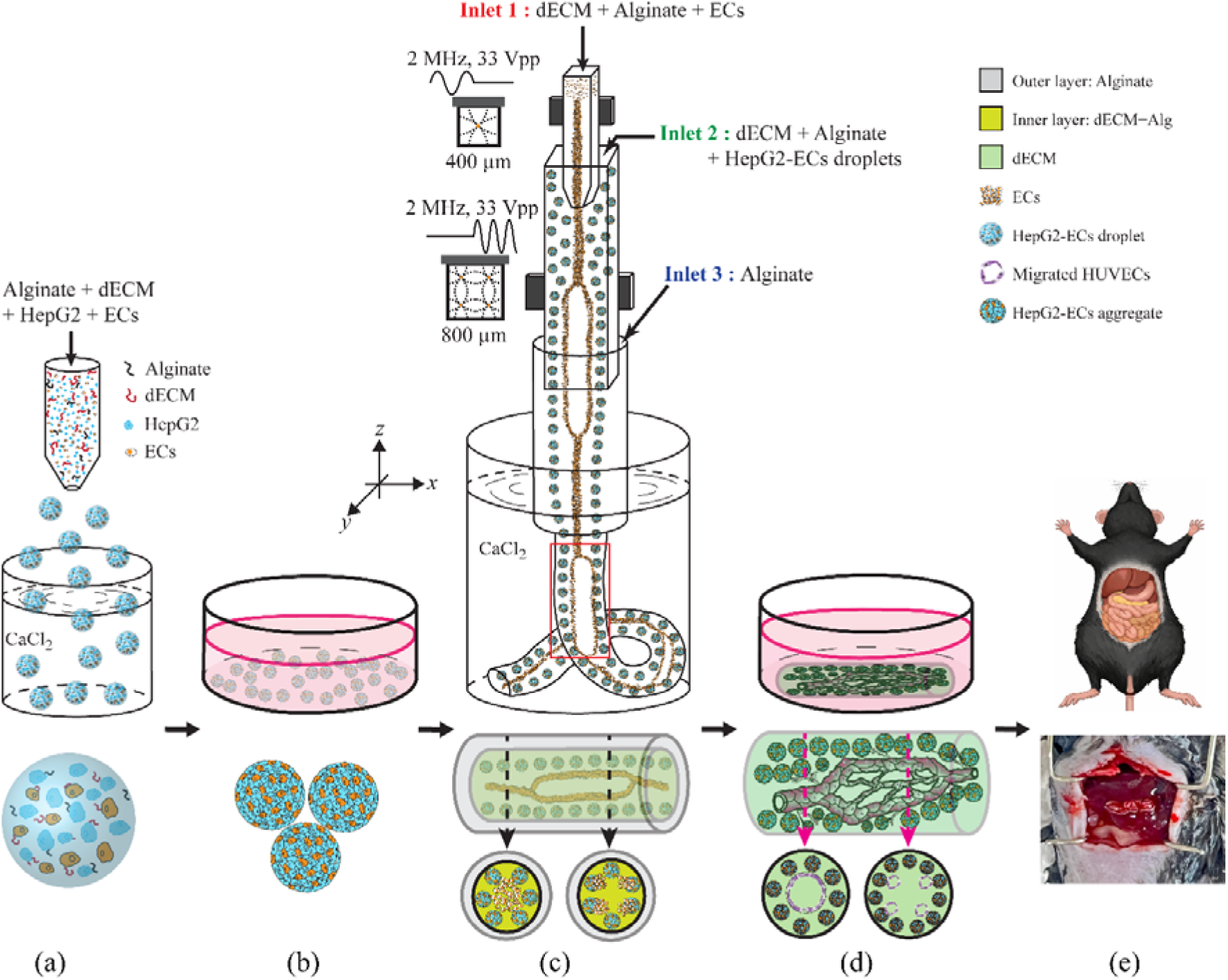
Schematic illustration of the artificial liver fabrication process using USW and HepG2–HUVEC aggregates. (a) Generation of the HepG2–HUVEC aggregates by spraying a cell-laden alginate/dECM suspension into a CaCl_2_ bath. (b) Fragmentation of the cell aggregates treatment following four days incubation. (c) Extrusion of the liver scaffold under ultrasound-wave patterning, generating spatially distinct regions of single-stream and four-stream HUVEC alignment by selective transducer actuation. (d) One-day culture of the fabricated liver scaffold without alginate lyase, followed by seven-day culture in the presence of alginate lyase to promote microvascular network formation.

### 2.2. Hydrogel preparation

Liver dECM powder (350 mg) and pepsin (200 mg; Sigma-Aldrich, St. Louis, MO, U.S.A.) were dissolved in a mixture of 5 mL acetic acid (Sigma-Aldrich, St. Louis, MO, U.S.A.) and deionized (DI) water to a final volume of 100 mL, yielding a dECM concentration of 35 mg/mL. The solution was stirred at room temperature (RT) for 72–96 h, and subsequently centrifuged at 3,000 rpm for 15 min. The supernatant was carefully transferred into separate conical tubes, and the undissolved residue was discarded. The tubes containing the solubilized dECM were stored at 2–8 °C.

For neutralization and dilution, the 35 mg/mL dECM stock solution was mixed with 10 M and 1 M NaOH (Sigma-Aldrich, St. Louis, MO, U.S.A.) and 10× PBS (10% of the total volume) to adjust the pH in the range of 7.2–7.5 and to yield a final dECM concentration of 25 mg/mL. The precursor solution used for acoustic patterning was then prepared by combining sodium alginate (0.5% w/v; MERCK, Madison, NJ, U.S.A.) and the neutralized dECM (25 mg/mL) at a volume ratio of 1:9. All procedures were performed on ice to prevent premature gelation.

### 2.3. Cell culture

HepG2 cells and the human umbilical vein endothelial cell line EA.hy926 were purchased from ATCC (Manassas, VA, U.S.A.). Culture medium consisted of Dulbecco’s Modified Eagle Medium (DMEM; Gibco, U.S.A.) supplemented with 10% fetal bovine serum (FBS; Gibco, U.S.A.) and 1% penicillin/streptomycin (Sigma-Aldrich, St. Louis, MO, U.S.A.). The cells were seeded onto polystyrene tissue culture dishes and maintained at 37 °C in a humidified atmosphere containing 5% CO. Cell growth was monitored using an IX53 inverted microscope (Olympus, Japan).

### 2.4. HepG2–EA.hy926 cell aggregates fabrication

HepG2 and EA.hy926 cells (3:1 ratio) were suspended in a mixture of dECM and sodium alginate at a volume ratio of 9:1 to fabricate HepG2–EA.hy926 cell (HG–EC) aggregates. This ratio was selected to maximize the supportive microenvironment provided by the dECM while preserving the structural integrity conferred by the sodium alginate. The resulting cell-laden precursor solution was then sprayed through a nozzle with an inner diameter of 75 ± 3 µm at a flow rate of 20 µL/min, and the droplets were collected in a bath of 0.1 M CaCl (MERCK, Madison, NJ, U.S.A.) for ionic crosslinking. The crosslinked microdroplets were subsequently cultured in the medium described above for four days, during which the cells self-assembled into HG–EC aggregates within each droplet (Fig. 2a). After this four-day culture, gentle pipetting alone was sufficient to fragment the HG-EC aggregates into smaller aggregates, without requiring alginate lyase treatment, owing to the low alginate content of the precursor solution.

### 2.5. Scaffold fabrication using USW

As shown in Fig. 2b, 1 mm-thick ultrasound transducers (MEGITT A/S, Kvistgaard, Denmark) were bonded to 400 µm and 800 µm square glass capillaries (VitroCom, NJ, U.S.A.), respectively. The device comprised three inlets. At inlet 1, a mixture of sodium alginate and liver dECM combined with HUVECs was introduced, while at inlet 2, the same alginate / liver dECM formulation combined with HepG2 aggregates was supplied. Inlet 3 was used to inject sodium alginate alone without any cells (1% w/v). The flow rates were set to 30 µL/min for inlets 1 and 2, and 300 µL/min for inlet 3. The outlet was immersed in a bath of 300 mM CaCl to induce ionic crosslinking. The fabricated scaffold was then incubated at 37 °C for at least 10 min to allow dECM gelation.

### 2.6. Microvascular network formation

The extruded artificial liver scaffolds were initially cultured in the conventional medium described above for 24 h at 37 °C in a humidified atmosphere containing 5% CO in order to preserve the integrity of the three-layered extruded structure. Alginate lyase (MERCK, Madison, NJ, U.S.A.) was then added to the culture medium at a final concentration of 0.1 U/mL to selectively degrade the calcium alginate component of the scaffolds, thereby generating pathways that enable the endothelial cells to migrate and integrate within the construct. The scaffolds were maintained in the enzyme-containing medium for a further seven days, with the medium refreshed every two days to sustain enzymatic activity and to prevent the accumulation of degradation byproducts that could otherwise impair cell migration or compromise scaffold integrity.

### 2.7. Cell staining

At 6 h and 24 h after scaffold fabrication, the cell-laden scaffolds were stained using a LIVE/DEAD assay. The staining solution consisted of 0.2% ethidium homodimer-1 (EthD-1; 2 mM in DMSO/H O, 1:4, v/v) and 0.05% calcein-acetoxymethyl (calcein-AM; 4 mM in anhydrous DMSO) from the LIVE/DEAD Viability/Cytotoxicity Kit for mammalian cells (Molecular Probes, Eugene, OR, U.S.A.), diluted in 1× phosphate-buffered saline (PBS; D8662-500ML, MERCK, Madison, NJ, U.S.A.). Before and after staining, the scaffolds were gently rinsed three times with 1× PBS (1–3 min per wash) to remove residual medium and unbound dye, and were subsequently imaged with a 4× objective on an Olympus BX53 upright fluorescence microscope (Olympus, Japan). Five representative images per sample were analyzed using ImageJ (Fiji, NIH, U.S.A.). For the quantification of cell viability, the fluorescence signals from labeled cells were separated into two channels: a red channel corresponding to dead cells (EthD-1) and a green channel corresponding to live cells (calcein-AM). The intensity of each channel was measured in ImageJ, and the percentage of viable cells was calculated as the ratio of green intensity to the sum of green and red intensities:

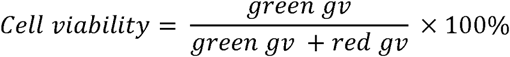

### 2.8. Proliferation assay

At 24 h of conventional culture and at days 3 and 7 following alginate lyase addition, the liver scaffolds were transferred to a 96-well plate containing 100 µL of fresh medium per well. Subsequently, 20 µL of CellTiter 96® AQueous One Solution Cell Proliferation Assay reagent (Promega, Madison, WI, U.S.A.), which contains a tetrazolium compound (3-(4,5-dimethylthiazol-2-yl)-5-(3-carboxymethoxyphenyl)-2-(4-sulfophenyl)-2H-tetrazolium; MTS) and an electron-coupling reagent (phenazine ethosulfate; PES), was added to each well, and the scaffolds were incubated at 37 °C for 4 h. The absorbance of the resulting formazan product was measured at 490 nm using a SpectraMax iD3 microplate reader (Molecular Devices, San Jose, CA, U.S.A.). All measurements were performed in six biological replicates under identical conditions.

To evaluate cell proliferation in the implanted scaffolds, the scaffolds were explanted on days 1, 7, 14, and 21 post-implantation. The explanted scaffolds were thoroughly rinsed with 1× PBS to remove residual blood cells, cut into small pieces (about 1 mm^3^), and processed according to the proliferation assay protocol described above. All experiments were performed in triplicate under identical conditions.

### 2.9. Urea production

To assess hepatocyte function during scaffold culturing, culture supernatants were collected at days 1, 3, 7 and analyzed. Synthesized urea was quantified using a urea assay kit (ab83362, Abcam, Cambridge, UK), according to the manufacturer’s instructions. Absorbance was measured 570 nm for urea using a SpectraMax iD3 microplate reader (Molecular Devices, San Jose, CA, U.S.A.).

For the animal experiments, blood samples were collected from the lateral tail veins of the mice on days 1, 7, 14, and 21 and immediately centrifuged at 2,000 × g for 10 min at room temperature to collect the blood plasma. After that, the urea concentrations were measured using the same urea assay kits described above, to evaluate hepatocyte function *in vivo* over time.

### 2.10. Immunofluorescence staining

The artificial liver scaffolds were fixed in 4% paraformaldehyde (Sigma-Aldrich, St. Louis, MO, U.S.A.) for 25 min at RT, permeabilized with 0.5% Triton X-100 (Sigma-Aldrich, St. Louis, MO, U.S.A.) for 5 min, and then blocked with 5% bovine serum albumin (BSA; Sigma-Aldrich, St. Louis, MO, U.S.A.) for 20 min at RT. The scaffolds were then incubated overnight at 4 °C with primary antibodies against CD31 (1:500; ab9498, Abcam, Cambridge, UK) and albumin (1:200; ab207327, Abcam, Cambridge, UK). After washing, the scaffolds were incubated for 120 min at RT with the corresponding secondary antibodies: goat anti-mouse IgG (H+L) highly cross-adsorbed Alexa Fluor® Plus 488 (1:1000; Thermo Scientific, Waltham, MA, U.S.A.) for CD31 detection, and goat anti-rabbit IgG (H+L) Alexa Fluor® 594 (1:1000; ab150080, Abcam, Cambridge, UK) for albumin detection. Nuclei were counterstained with 4′,6-diamidino-2-phenylindole (DAPI; NucBlue^®^ Fixed ReadyProbes™ Reagent, Thermo Scientific, Waltham, MA, U.S.A.) for 15 min prior to imaging. Between each step, the scaffolds were washed three times with 1× PBS buffer for 5 min per wash.

### 2.11. qPCR

For RNA extraction, the cell aggregates and scaffolds were lysed in 500 µL of TRIzol™ reagent (Gibco, U.S.A.) by repetitive pipetting. Chloroform (50 µL) was added to the lysate, followed by incubation for 3–5 min at room temperature. The samples were then centrifuged at 12,000 × *g* for 10 min at 4 °C. The colorless aqueous phase was carefully transferred to a gDNA Eliminator spin column from the RNeasy® Plus Mini Kit (Qiagen, Germany), and total RNA was purified according to the manufacturer’s instructions.

Total RNA was reverse-transcribed into cDNA using the High-Capacity cDNA Reverse Transcription Kit (Thermo Fisher Scientific, Waltham, MA, U.S.A.) according to the manufacturer’s instructions.

Quantitative PCR (qPCR) was performed for *AFP*, *CK18*, *CYP1A2*, *CYP2E1*, *CYP2D6*, and *CYP3A4* using Maxima SYBR Green/ROX qPCR Master Mix (Thermo Fisher Scientific, Waltham, MA, U.S.A.) with the primers (Bioneer, Daejeon, Republic of Korea) listed in Table 1, according to the manufacturer’s protocol. All qPCR reactions were carried out on a LightCycler® 480 Instrument II (Roche, Basel, Switzerland), and the resulting data were analyzed using LightCycler® 480 Software (Roche). Relative gene expression levels were calculated using the 2^(−ΔΔCT) method with β-actin as the reference gene (Duong et al., Biofabrication, 2020; Mori et al., The Prostate, 2008). Each qPCR reaction was performed in at least three technical replicates per sample.

**Table 1:**
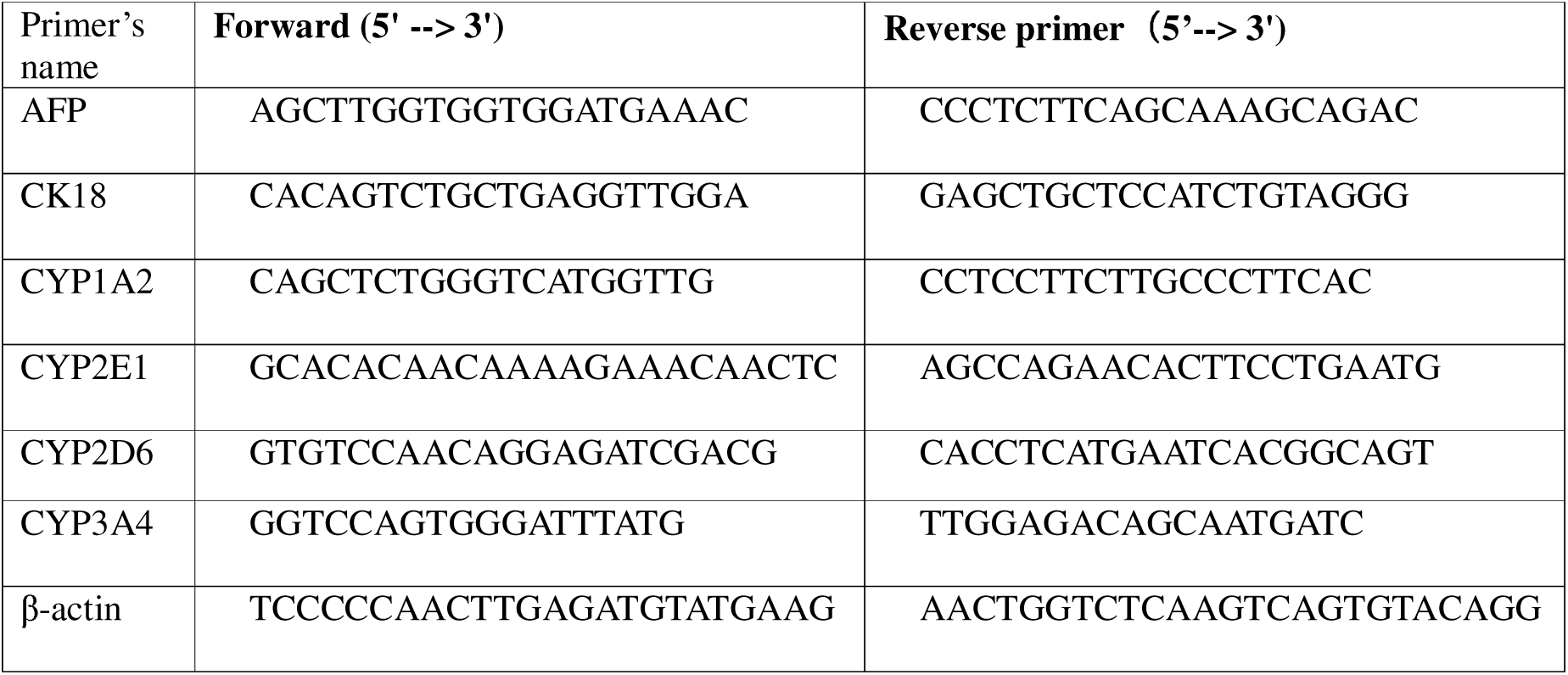
Primer’s consequence.

### 2.12. Implant surgery

Mice (male, 6–8 weeks old, 20–30 g; C57BL/6, Hana Biotech, Republic of Korea) were anesthetized by intraperitoneal injection of 2,2,2-tribromoethanol (∼240 mg/kg; Sigma-Aldrich, St. Louis, MO, U.S.A.). The abdominal hair was removed using Nair™ depilatory gel (Church & Dwight UK Ltd., Kent, U.K.), and the surgical site was disinfected with 10% povidone-iodine solution (Forson, Chungcheongnam, Republic of Korea). The abdominal skin and underlying muscle layer were incised using surgical scissors and fine forceps, after which a small portion of the liver was partially resected to accommodate the scaffold. After eight days of culture, the matured scaffold (∼2 cm in length) was placed into the resection site and secured with 8-0 black silk sutures (Ailee, Busan, Republic of Korea). The skin incision was then closed with 4-0 black silk sutures (Ailee, Busan, Republic of Korea). Following surgery, the animals were placed on a heating pad and monitored for 3–4 h until full recovery from anesthesia. Each mouse was housed individually, and signs of inflammation, distress, or wound complications were assessed visually throughout the postoperative period. At days 1, 7, 14, and 21 post-implantation (Fig. 3), the mice were re-anesthetized and the implanted scaffolds were harvested, after which the animals were euthanized by cervical dislocation. All surgical instruments were autoclaved prior to use. All animal care and procedures were conducted according to the protocols and guidelines approved by the University of Ulsan Animal Care and Use Committee (UOUACUC) (Permit number: GIG-23-020).

**Figure 3.**
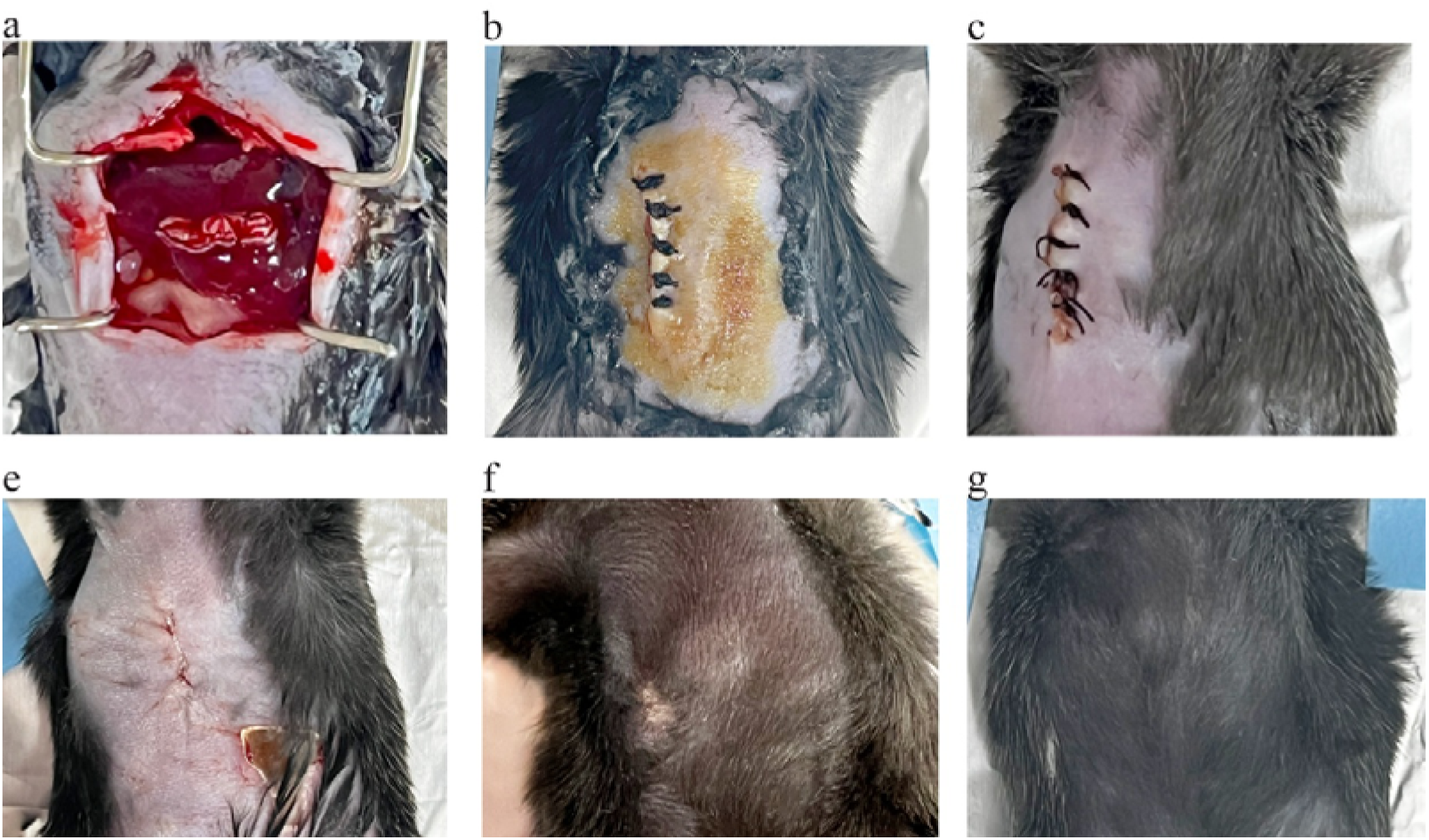
Implantation of the matured artificial liver scaffold into a mouse. (a) The matured scaffold was positioned within the resection cavity created by removing a corresponding volume of liver tissue. (b) The abdominal incision was closed with sutures immediately after implantation. The implanted regions are shown at (c) 1 day, (d) 7 days, (e) 14 days, and (f) 21 days post-implantation.

### 2.13. Imaging and statistical analysis

The stained scaffolds were observed under an IX53 inverted fluorescence microscope (Olympus, Tokyo, Japan), and images were acquired using CellSens software (Olympus, Tokyo, Japan). Three-dimensional images were obtained using a laser-scanning confocal microscope (FLUOVIEW FV1200, Olympus, Tokyo, Japan). Image processing and quantitative analysis were performed using ImageJ software (NIH, U.S.A.). All data are presented as mean ± standard deviation (SD).

## 3. Results

### 3.1. HG-EC aggregates culturing

Culturing hepatocytes as three-dimensional (3D) aggregates can enhance their functional performance by promoting cell-cell contact within a tissue-like microenvironment. Compared with conventional two-dimensional (2D) monolayer cultures, hepatocyte aggregates more closely resemble native liver tissue and can be regarded as micro-tissues. Moreover, such aggregates can serve as functional building blocks for the bottom-up assembly of larger and hierarchically organized constructs. Although HepG2 cells, as a hepatocellular carcinoma cell line, do not fully recapitulate the *in vivo* phenotype of primary hepatocytes, they are easy to handle, robust, and readily form aggregates while retaining several hepatocyte-specific functions.

The aggregation process was monitored at three different cell seeding densities: 1 × 10^6^, 3 × 10^6^, and 5 × 10^6^ cells/mL. Within the sprayed droplets, cells migrated and self-assembled into multiple aggregates over time (Fig. 4a, b). After four days of culture, the mean diameters increased from 23.3 ± 6.2 µm to 41.1 ± 11.7 µm at 1 × 10^6^ cells/mL (76.4% increase), from 23.4 ± 7.1 µm to 50.3 ± 16.5 µm at 3 × 10^6^ cells/mL (115.0% increase), and from 28.9 ± 6.6 µm to 77.5 ± 19.1 µm at 5 × 10^6^ cells/mL (168.2% increase) (Fig. 4d). The coefficient of variation (CV; defined as the standard deviation divided by the mean) increased only slightly during the same period, from 26.6% to 28.5% at 1 × 10 cells/mL (Δ = +1.9 percentage points), from 30.3% to 32.8% at 3 × 10 cells/mL (Δ = +2.5 percentage points), and from 22.8% to 24.6% at 5 × 10 cells/mL (Δ = +1.8 percentage points), indicating that aggregate size uniformity was largely preserved during culture.

**Figure 4.**
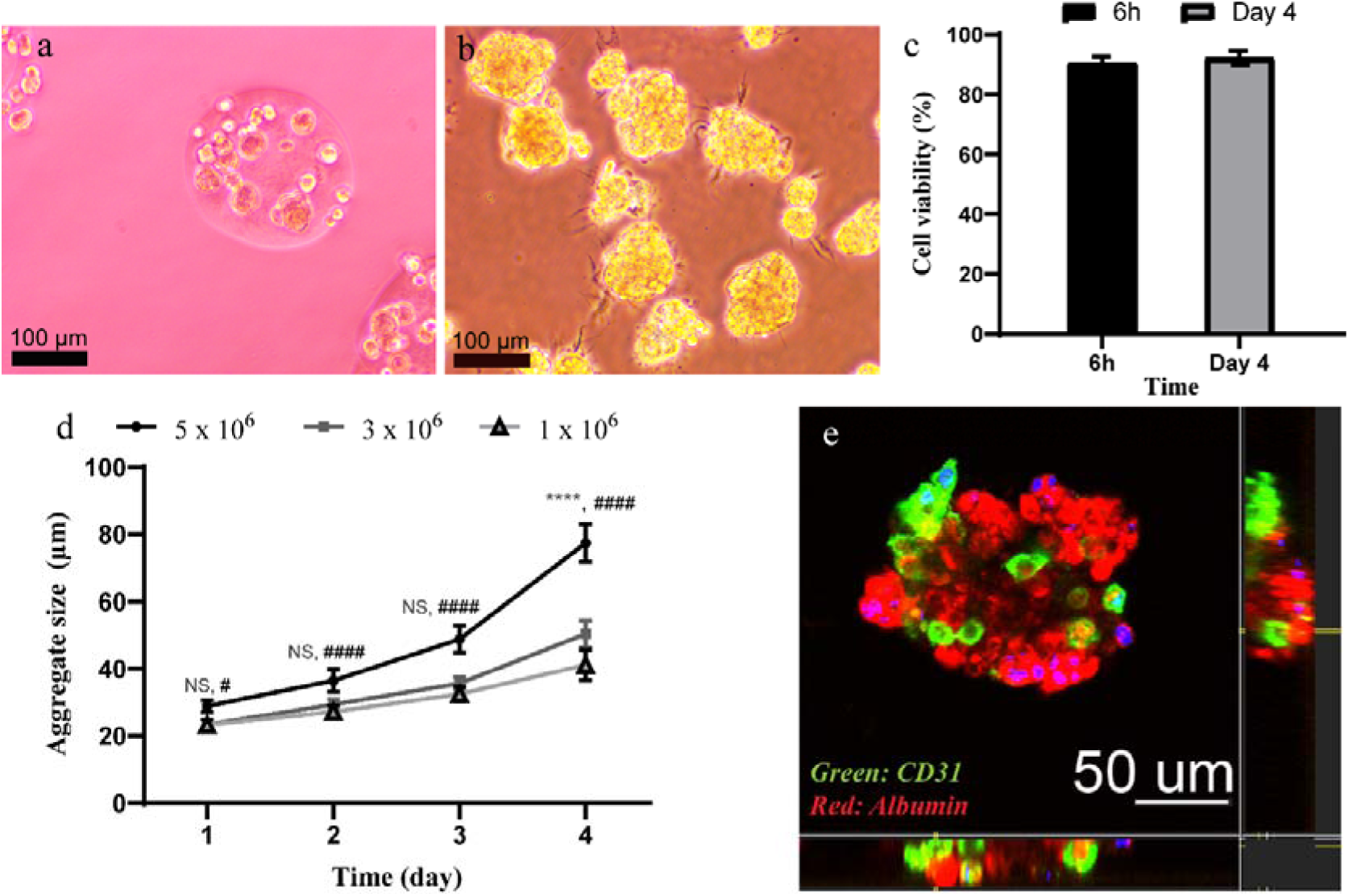
Fabrication and culture of HG–EC aggregates. Brightfield images of the HG–EC aggregates on (a) day 0 and (b) day 4. (c) Cell viability at 6 h and on day 4 after fabrication (n = 5). (d) Relationship between cell seeding density and aggregate size after four days of culture (# *p* < 0.05; ****, #### *p* < 0.0001 compared with the 1 × 10 cells/mL aggregates; n = 100). (e) Confocal fluorescence image of the HG–EC aggregates on day 4: CD31 (green), albumin (red), and nuclei stained with DAPI (blue).

Furthermore, LIVE/DEAD staining demonstrated excellent cell viability in all three seeding densities, with HG-EC aggregate viability exceeding 90% on day 4 of culture (Fig. 4c). The aggregates exhibited a compact spherical morphology composed of densely packed HepG2 and EA.hy926 cells (Fig. 4e).

### 3.2. Artificial liver scaffolds formation with branched structures of ECs

To fabricate three-dimensional vascularized liver scaffolds, the cultured HG–EC aggregates were mixed with single endothelial cells and subjected to USW patterning. Owing to their positive acoustic contrast factor, the single endothelial cells were focused toward the pressure nodes, forming alternating single-line and four-line patterns within the hydrogel (Fig. 5), thereby generating branched vascular structures. In contrast, the HG–EC aggregates, which exhibit a negative contrast factor, accumulated at the pressure antinodes located along the capillary sidewalls. These results demonstrate that the proposed USW-mediated patterning approach can fabricate a fundamental structural unit of liver tissue, comprising branched vascular networks surrounded by hepatocyte clusters.

**Figure 4.**
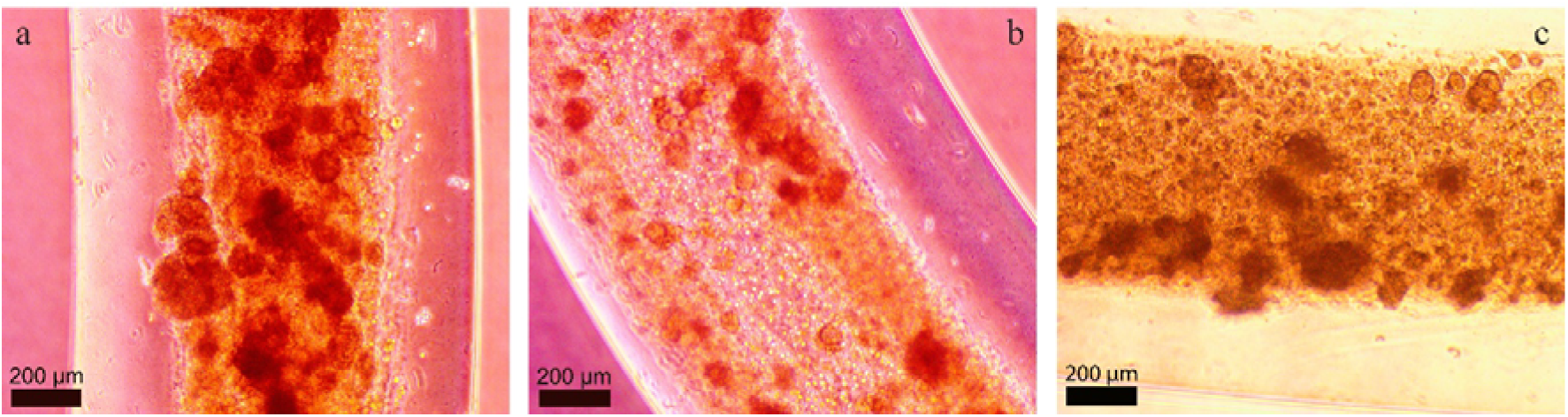
Artificial liver scaffolds patterned by USW. (a) Endothelial cells aligned as a single stream by 2 MHz ultrasound actuation in a 400 μm square glass capillary. (b) Endothelial cells aligned into four streams by 2 MHz ultrasound in an 800 μm square glass capillary. (c) Control scaffold fabricated without ultrasound actuation showing randomly distributed cells.

**Figure 5.**
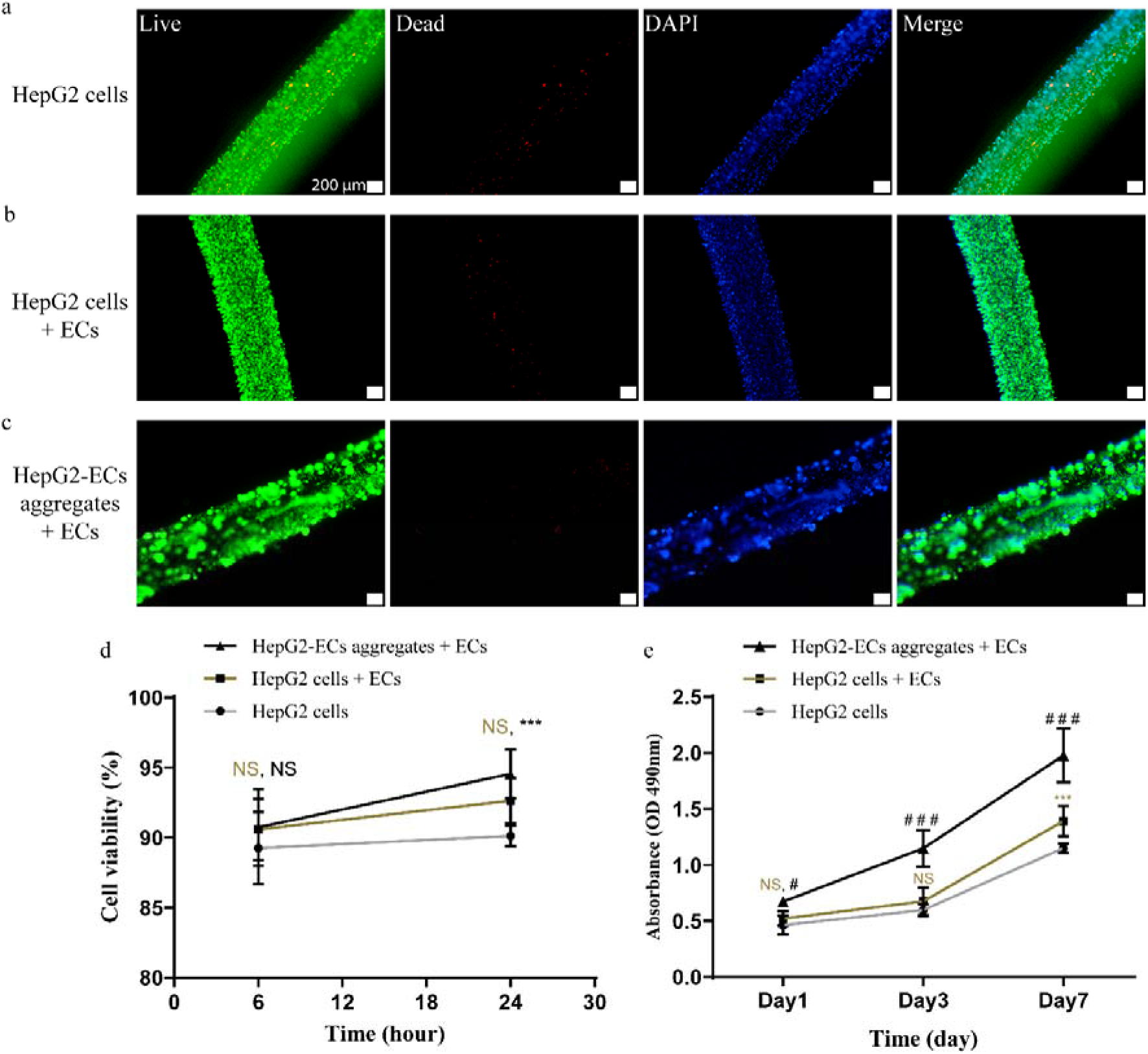
Evaluation of cell viability and proliferation in the fabricated scaffolds. (a–c) LIVE/DEAD fluorescence images of scaffolds containing (a) HepG2 cells only, (b) HepG2 cells and endothelial cells, and (c) HG-EC aggregates and endothelial cells, acquired at 24 h after extrusion. (d) Cell viability at 6 h and 24 h after extrusion. (e) Cell proliferation evaluated by the CellTiter 96^®^ AQueous One Solution Cell Proliferation Assay over seven days of culture following alginate lyase addition. Live cells are shown in green (calcein-AM), dead cells in red (ethidium homodimer-1, EthD-1), and nuclei in blue (DAPI). Data are presented as mean ± SD (n = 6; #p < 0.05; ***, ### p < 0.001; n.s., not significant; compared with the HepG2-only scaffold).

### 3.3. Microvascular network formation in the artificial liver scaffolds and perfusibility function

The cell viability in the two scaffolds including endothelial cells increased and reached approximately 90% within one day after extrusion (Fig. 6d). The proliferation profiles of the generated scaffolds over seven days following alginate lyase treatment are shown in Fig. 6e. The scaffold containing HG-EC aggregates and endothelial cells exhibited markedly faster proliferation than other scaffolds, with an approximately two-fold higher proliferation rate compared with the HepG2-only scaffold.

**Figure 6.**
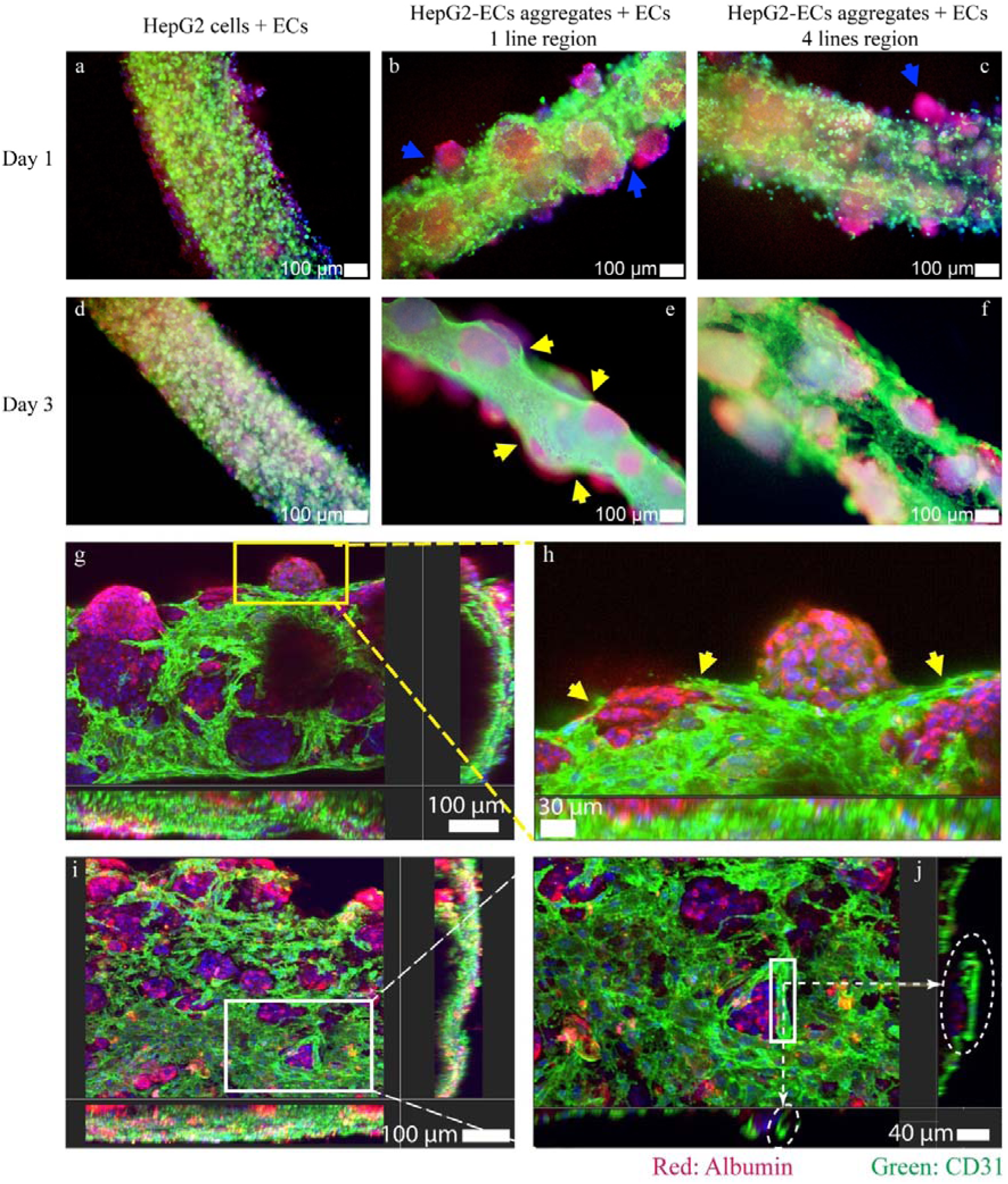
Immunofluorescence images of the fabricated scaffolds. **(a–c)** Confocal images of **(a)** the HepG2 + EC scaffold, **(b)** the single-line region of the HG-EC + EC scaffold, and **(c)** the four-line region of the HG-EC + EC scaffold on day 1 of alginate lyase treatment. **(d–f)** Corresponding images of the same regions on day 3. **(g, h)** Hollow channels formed in the single-line region and **(i, j)** branched lumen structures in the four-line region after three days of culture in the presence of alginate lyase. CD31 is shown in green (endothelial cell marker), albumin in red (hepatocyte marker), and nuclei in blue (DAPI). Blue arrows indicate aggregates without endothelial migration, and yellow arrows indicate endothelial cells migrating into the aggregates; white arrows indicate endothelial lumen formation.

Endothelial cell spreading within the artificial liver scaffolds was visualized by immunofluorescence staining for CD31 (green), a biomarker of platelet endothelial cell adhesion molecule, and albumin (red), a marker of HepG2 cells (Fig. 7). On day 1 of alginate lyase treatment, a subset of the endothelial cells retained their initial spherical morphology within the hydrogel. By day 3 of alginate lyase treatment, the scaffold containing HG-EC aggregates and endothelial cells (Fig. 7e, f) exhibited robust cell growth compared with that containing the HepG2 cells and endothelial cells (Fig. 7d). Confocal fluorescence imaging of CD31 and albumin further revealed lumen formation by endothelial cells in the single-line regions (Fig. 7g, h) and the maturation of interconnected vascular networks in the four-line regions (Fig. 7i, j). The endothelial cells also migrated into and spread within the HG-EC aggregates on day 3 (the yellow arrows in Fig. 7e, g, h), not like those on day 1 (the blue arrows in Fig 7b, c). Collectively, these results demonstrate that endothelial cells support and enhance the growth of hepatocytes in the extruded artificial liver scaffolds.

**Figure 7.**
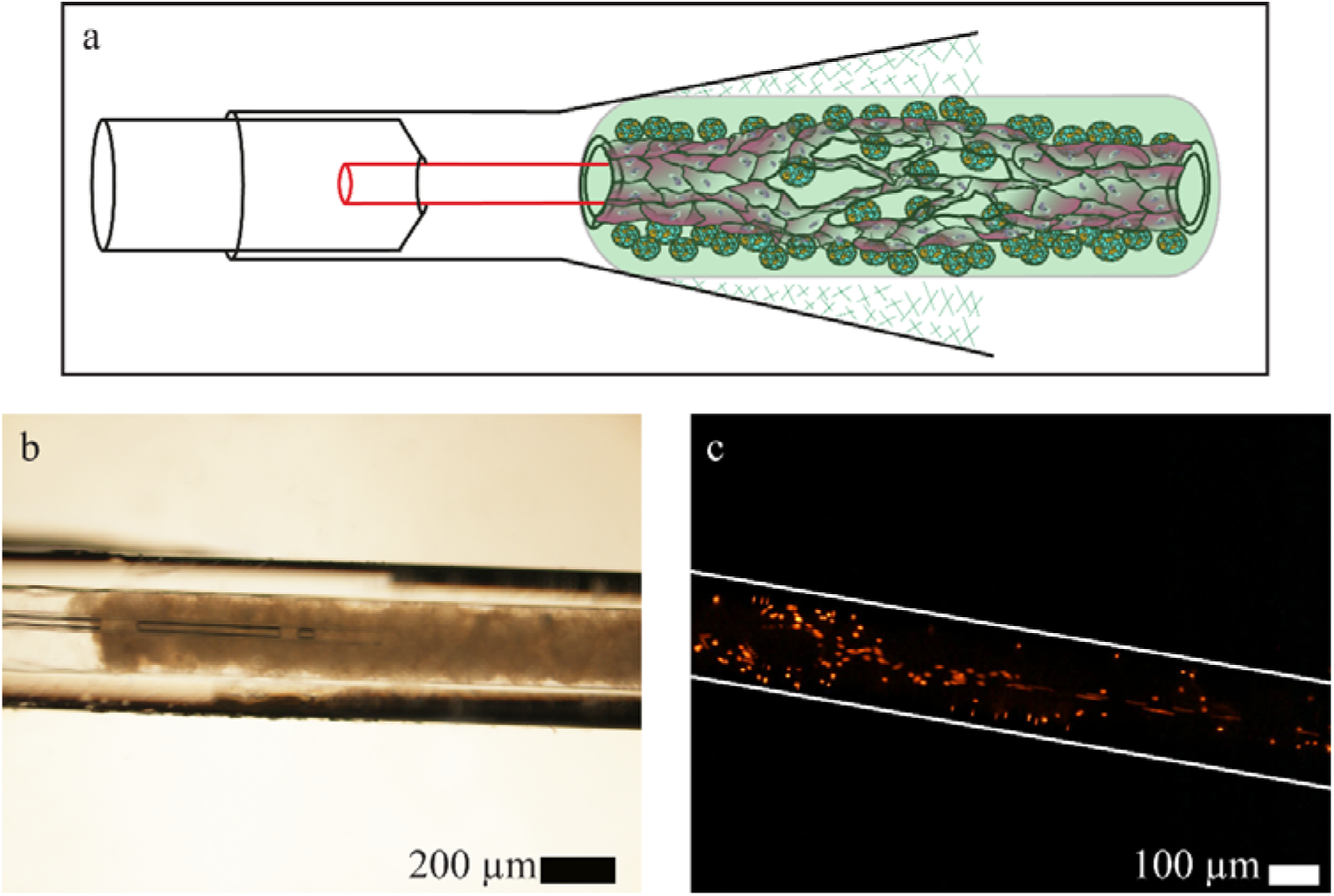
Perfusion assessment of the vascularized artificial liver scaffolds. (a) Schematic illustration of the connector setup used to interface the perfusion system with the liver scaffold. (b) Brightfield image of the scaffold-connector interface prior to perfusion. (c) Fluorescence image of the scaffold during perfusion of fluorescent microparticles (5 µm in diameter), shown as red.

Perfusion is a critical functional property of vascular systems, as it is essential for the establishment of stable blood or nutrient circulation within organs and tissues (Kim et al., Lab Chip, 2007; Sosa et al., Clin Hemorheol Microcirc, 2014). To assess the perfusability of the fabricated scaffold, fluorescent microparticles were perfused through the cultured scaffolds using a custom-built connector (Fig. 8a) (Le et al., Biofabrication, 2023). As shown in Fig. 8b and Supplementary Video 1, 5 µm fluorescent microparticles flowed freely through the endothelial channels. These results indicate that the engineered vascular networks possess an intact endothelial lining, as supported by biomarker expression, and are capable of effectively confining perfused fluids within the lumen.

**Figure 8.**
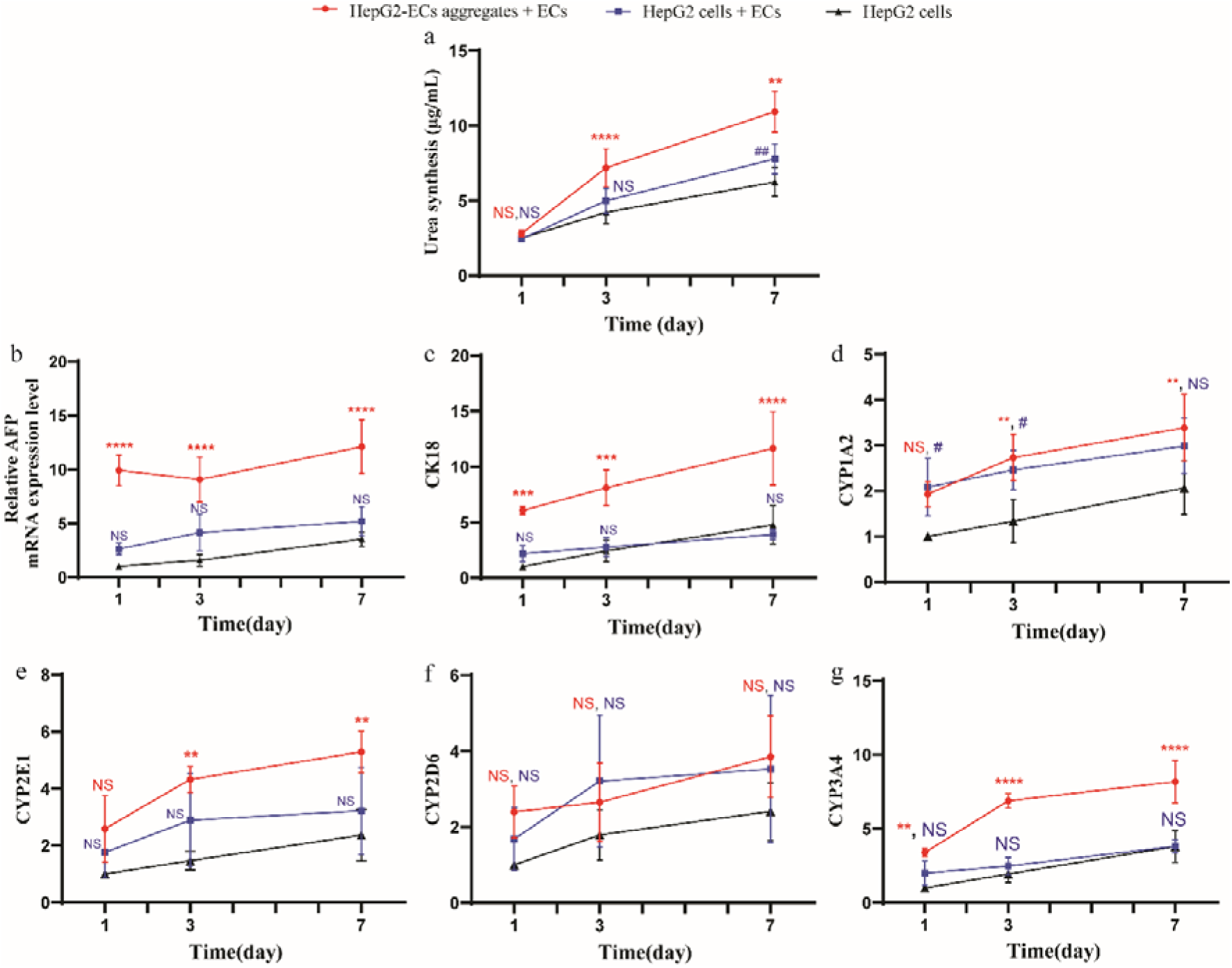
Evaluation of hepatocyte-specific functional expression in the artificial liver scaffolds. (a) Urea production, measured as markers of hepatocyte activity, on days 1, 3, and 7 of culture after adding alginate lyase. (b-g) Relative gene expression of (b) *AFP*, (c) *CK18*, (d) *CYP1A2*, (e) *CYP2E1*, (f) *CYP2D6*, and (g) *CYP3A4*, quantified by qPCR and normalized to β*-actin*. Data are presented as mean ± SD from three independent experiments (n = 3; \**p* < 0.05, \*\**p* < 0.01, \*\*\**p* < 0.001, \*\*\*\**p* < 0.0001, compared with the HepG2-only scaffold; NS, not significant).

### 3.4. HG-EC aggregates with vascularized networks enhanced liver-specific functional expression

The liver performs a variety of essential functions, including protein synthesis and metabolism. The urea production, an indicator of hepatic metabolic activity, is widely used as functional markers of hepatocytes (Lee et al., Biofabrication, 2016; Su et al., Genet Mol Res, 2016). Culture supernatants were collected on days 1, 3, and 7 of culture in the presence of alginate lyase and subsequently analyzed. Urea production in the HG-EC + EC scaffold increased from 2.50 ± 0.2 µg/mL on day 1 to 10.96 ± 1.4 µg/mL on day 7, 1.7-fold higher than that of the HepG2-only scaffold (6.25 ± 1.0 µg/mL on day 7), as shown in Fig. 9a. The expression of AFP (α-fetoprotein), a marker of fetal hepatocytes, and CK18 (cytokeratin 18), a major intermediate filament protein expressed in hepatocytes, was analyzed together with cytochrome P450 (CYP) enzymes (*CYP1A2*, *CYP2E1*, *CYP2D6*, and *CYP3A4*), which are essential for hepatic metabolic activity (Acun et al., Bioengineering, 2022; Jin et al., iScience, 2024; Shrimali et al., Drug Metab Dispos, 2025). The expression levels of *AFP* and *CK18* were markedly upregulated in the HG-EC + EC scaffold compared with the HepG2-only scaffold (Fig. 9b, c). Similarly, the expression of the metabolic enzyme genes *CYP1A2*, *CYP2E1*, *CYP2D6*, and *CYP3A4* increased over seven days of culture in the presence of alginate lyase (Fig. 9d-g), with *CYP1A2* and *CYP3A4* showing 2.7- and 6.2-fold increases, respectively, in the HG-EC + EC scaffold. Collectively, these results demonstrate that the incorporation of HG-EC aggregates with vascularized networks into the artificial liver scaffolds enhances hepatocyte-specific functional expression.

**Figure 9.**
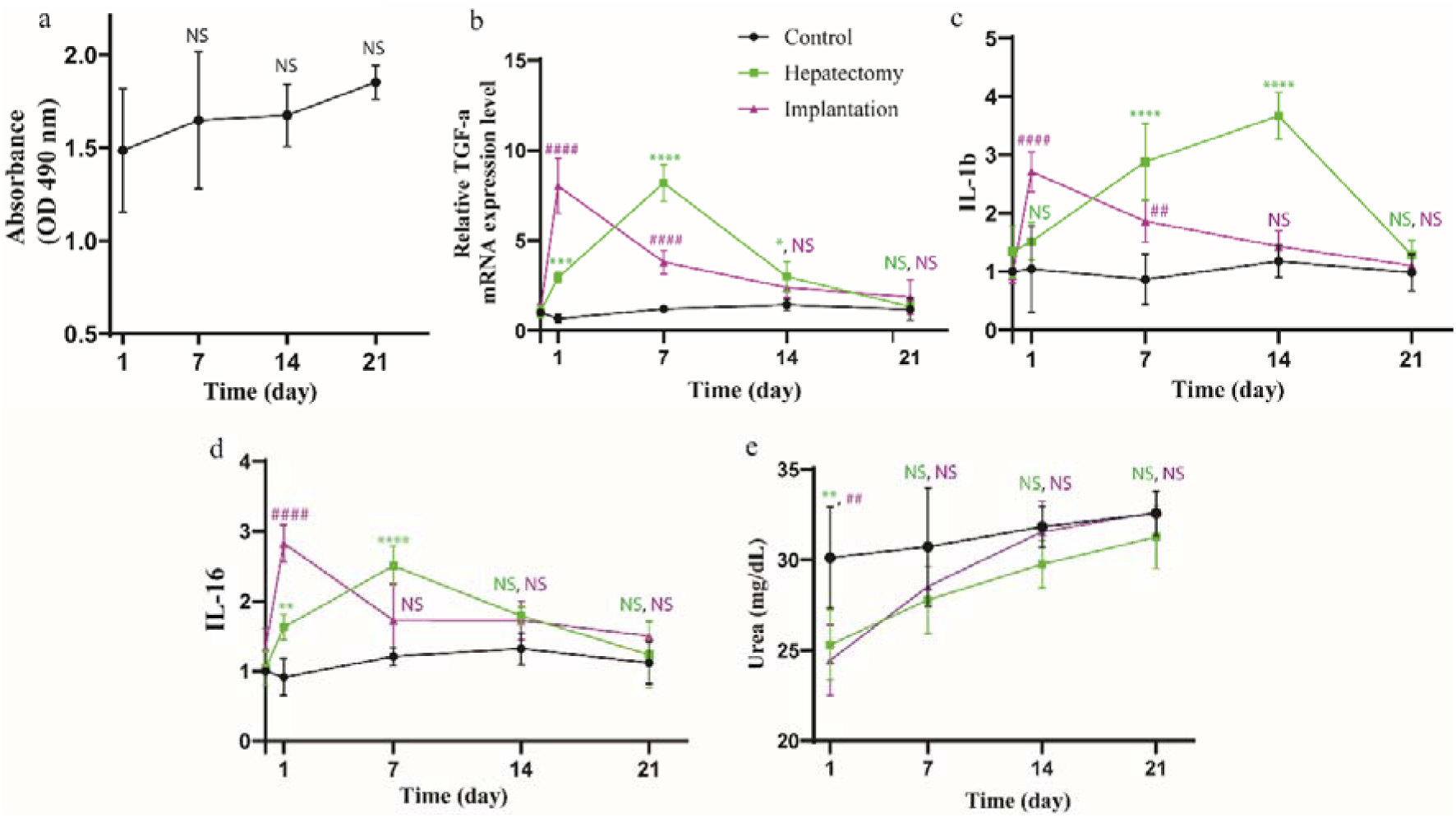
*In vivo* evaluation of the implanted artificial liver scaffolds. (a) Proliferation of the explanted scaffolds assessed by the CellTiter 96® AQueous One Solution Cell Proliferation Assay. Relative expression of (b) *TGF-*α, (c) *IL-1*β, and (d) *IL-16* in the surrounding liver tissue at different time points after implantation, quantified by qPCR and normalized to β*-actin*. (e) Time courses of urea production measured in the recipient mice following scaffold implantation. Data are presented as mean ± SD (n = 3 mice; \*\**p* < 0.01, \*\*\**p* < 0.001, \*\*\*\**p* < 0.0001, compared with the hepatectomy mouse; ##*p* < 0.01, ###*p* < 0.001, ####*p* < 0.0001, compared with the implantation mouse; NS, not significant).

### 3.5. In vivo experiment

The implantability of the artificial liver scaffolds was assessed by implanting them into the mouse liver (Fig. 2a). During the 21-day observation period, the implanted mice exhibited no signs of swelling, fever, or infection, and maintained normal vitality and behavior. The graft site showed rapid recovery, with complete skin healing by day 7 and hair regrowth by day 21 (Fig. 2b–f).

To evaluate whether the scaffolds retained their proliferative capacity following hepatic implantation, the explanted scaffolds were subjected to proliferation assays. The scaffolds exhibited sustained proliferation and remained viable throughout the 21-day post-implantation period (Fig. 10a). To assess the host response to the implanted scaffolds, cytokine profiling was performed for *TGF-*α (transforming growth factor-α), which is associated with proliferative and regenerative signaling, and for the pro-inflammatory and immune-regulatory cytokines *IL-1*β and *IL-16* (Kucsera et al., Scientific Reports, 2023; Sandgren et al., Mol Cell Biol, 1993; Santoni-Rugiu et al., Am J Pathol, 1996; Wen et al., Nature Communications, 2025). All three cytokines were markedly upregulated in both injury mice (the scaffold-implantation mouse and the partial-hepatectomy mouse) relative to the sham control mouse, which remained near baseline throughout the observation period; however, the two mice differed in the kinetics of this response. In the scaffold-implantation mouse, TGF-α expression peaked on day 1 at 8.0-fold and declined progressively to 1.9-fold by day 21, whereas in the partial-hepatectomy mouse it peaked later, on day 7, at 8.2-fold (Fig. 10b). The pro-inflammatory cytokines followed the same pattern: IL-1β and IL-16 peaked on day 1 in the implantation mouse (2.7- and 2.8-fold, respectively), whereas in the hepatectomy mouse IL-1β peaked on day 14 (3.7-fold) and IL-16 on day 7 (2.5-fold) (Fig. 10c, d). By day 21, all cytokine levels had returned toward baseline in both mice.

**Figure 10.**
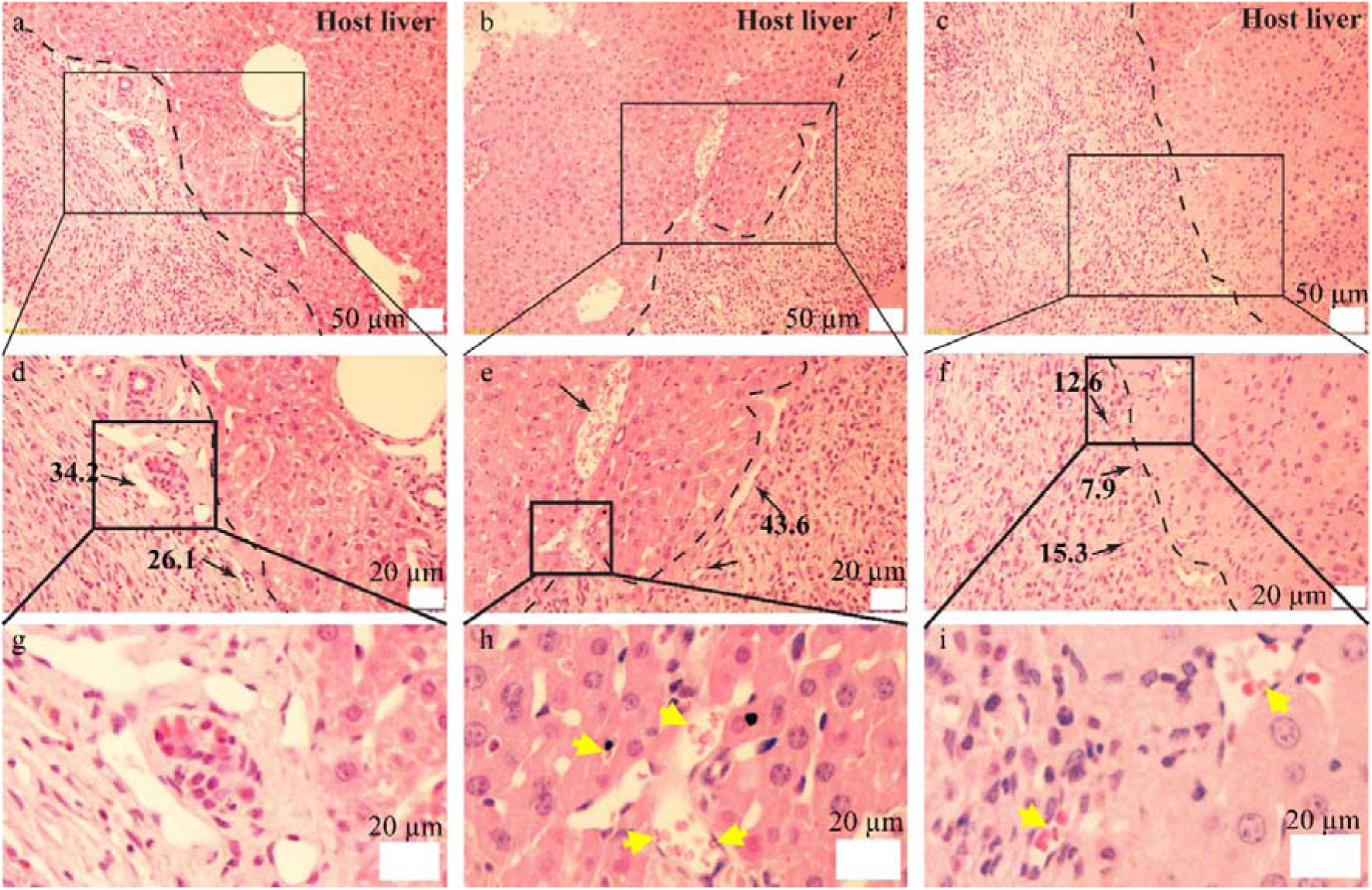
Hematoxylin and eosin (H&E) staining of the artificial liver scaffolds implanted into the mouse liver at (a, d) day 7, (b, e) day 14, and (c, f) day 21 post-implantation. Black arrows indicate microvessels containing red blood cells; yellow arrows indicate red blood cells. Numbers denote the diameters of the vascular lumina in micrometers (µm). Pink: extracellular matrix; purple: nuclei; bright red: red blood cells.

Urea production decreased significantly in both injury mice on day 1 compared with the sham control mouse, reaching 24.46 ± 1.96 mg/dL in the scaffold-implantation mouse and 25.31 ± 1.95 mg/dL in the hepatectomy mouse versus 30.12 ± 2.79 mg/dL in the sham control mouse (Fig. 10e). Thereafter, urea production recovered progressively in both mice, and from day 7 onward no significant difference from the sham control mouse was detected. By day 21, urea production in the scaffold-implantation mouse (32.64 ± 1.20 mg/dL) was comparable to that of the sham control mouse (32.58 ± 1.24 mg/dL), indicating that hepatic metabolic function was restored following implantation. The resulting values of the scaffold-implantation mouse remained within the physiological range reported for mice (Nguyen et al., Journal of Functional Biomaterials, 2025; Otto et al., J Am Assoc Lab Anim Sci, 2016).

Histological analyses of liver tissue collected on days 7, 14, and 21 revealed progressive integration between the scaffold and the host liver, with increasing invasion of host tissue into the scaffold over time, as observed by hematoxylin and eosin (H&E) staining (Fig. 11) and Masson’s trichrome staining (Fig. 12). Red blood cells were detected within progressively smaller lumina, with the smallest blood-containing lumen decreasing in diameter from 26.1 µm on day 7 to 7.9 µm by day 21, indicating the maturation of initially empty tubular structures into functional vessels and potential anastomosis with the host vasculature. The luminal contents were identified as red blood cells based on their anucleate, uniformly sized, biconcave morphology, in contrast to the clearly stained nuclei of the surrounding hepatocytes and lumen-lining cells in the same sections (the yellow arrows in Fig. 11h, i and 12h, i). In addition, Masson’s trichrome staining revealed a marked increase in collagen fiber deposition within the grafted tissue by day 21, indicating extracellular matrix remodeling and successful integration of the scaffold with the host tissue.

**Figure 11.**
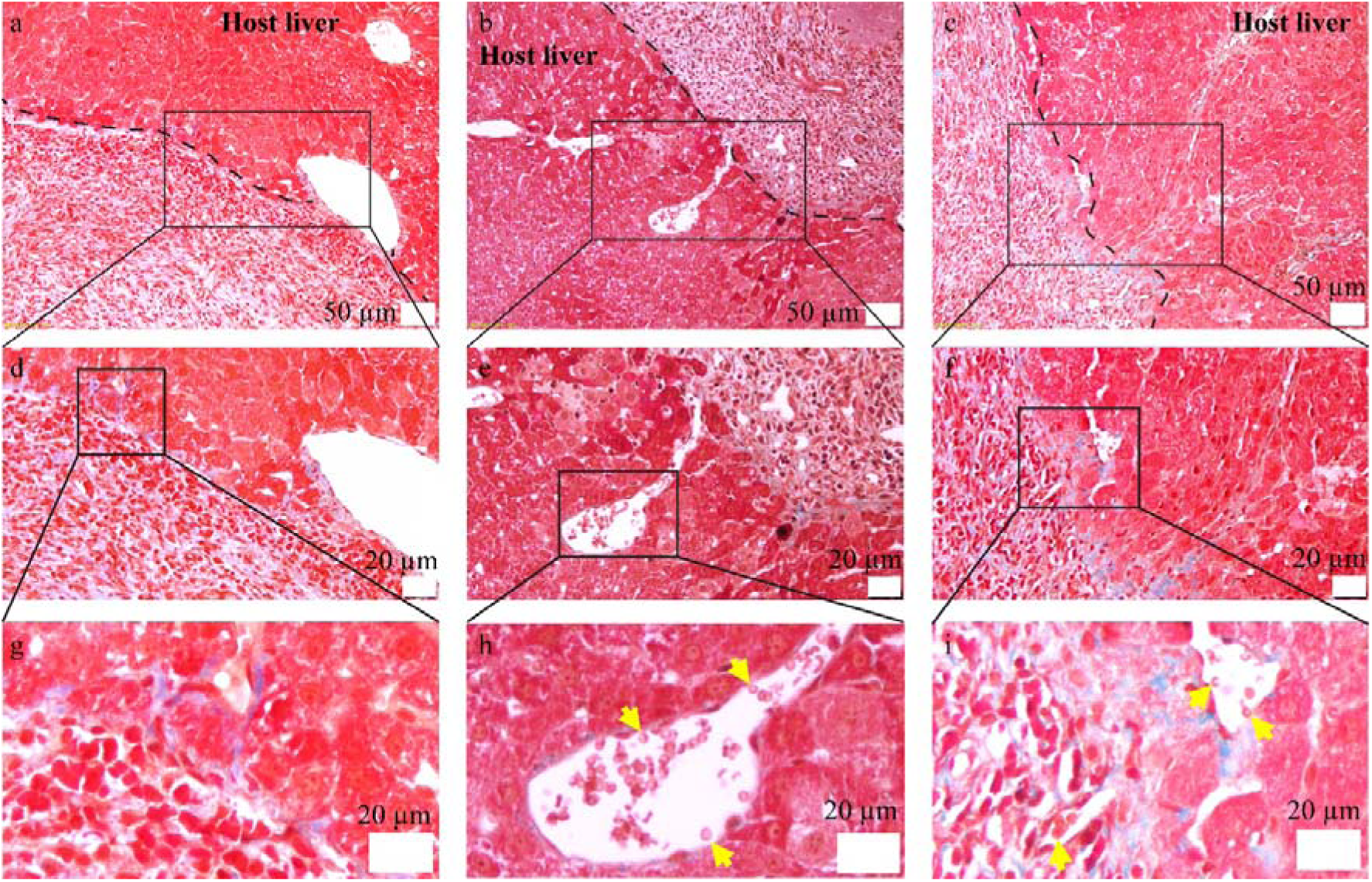
Masson’s trichrome staining of the artificial liver scaffolds implanted into the mouse liver at (a, d) day 7, (b, e) day 14, and (c, f) day 21 post-implantation. Microvessels are identified as lumen structures containing red blood cells. Yellow arrows indicate red blood cells. Red: cytoplasm; dark red: nuclei; bright red: red blood cells; light blue: collagen fibers and extracellular matrix.

## 4. Discussion

In the present study, we developed a strategy that integrates HG-EC aggregates with acoustically patterned endothelial networks to closely mimic the key structural and functional features of native liver tissue. The formation of HG-EC aggregates represents a critical step toward recreating the three-dimensional microenvironment of the liver. Compared with two-dimensional cultures, hepatocyte aggregation enhances cell–cell interactions and supports hepatic phenotypic stability (Liu et al., J Cell Biochem, 2007; Tutty et al., Drug Delivery and Translational Research, 2022). We observed that increasing the initial cell density during aggregation markedly influenced aggregate growth. Mean diameters increased by 76.4%, 115.0%, and 168.2% over four days at seeding densities of 1 × 10, 3 × 10, and 5 × 10 cells/mL, respectively, with the highest density yielding the largest aggregates (77.5 ± 19.1 µm; Fig. 4). This trend is consistent with the notion that dense three-dimensional assemblies promote hepatocyte maturation. Notably, the coefficient of variation in aggregate diameter increased by no more than 2.5 percentage points across all densities over the same period, and viability remained above 90% on day 4, indicating that aggregate size uniformity and cell survival were well preserved despite the substantial increase in size. This reproducibility supports the use of hepatocyte-endothelial aggregates as building blocks that preserve high local cellularity within the assembled construct. These aggregates function as liver-like microtissues and provide modular building blocks for the assembly of larger, hierarchically organized constructs (Jakab et al., Biofabrication, 2010; Lu et al., Biomaterials, 2025).

Beyond hepatocyte aggregation, the spatial organization of endothelial cells within the scaffold plays a pivotal role in supporting scaffold functionality. Using USW patterning, single endothelial cells were arranged into branched architectures within the hydrogel, while the hepatocyte aggregates localized to the surrounding regions. Critically, unlike bioprinting approaches that rely on sacrificial templates, coaxial nozzles, or ultraviolet crosslinking (Banaeiyan et al., Biofabrication, 2017; Wang et al., Frontiers in Bioengineering and Biotechnology, 2023; Wu et al., Scientific Reports, 2020), our strategy organizes both the vascular and parenchymal compartments concurrently within a single extrusion step, avoiding potential ultraviolet-induced cytotoxicity and the need for sacrificial materials. The resulting liver-mimetic unit comprises branched vascular channels surrounded by hepatocyte clusters (Fig. 5), in which the transition from a single-line to a four-line configuration forms a simple branched hierarchy connecting a larger supplying channel to finer parallel microchannels. This architecture reconstitutes the basic microvascular-parenchymal module of the hepatic lobule, in which hepatocytes are intimately associated with a branched vascular network (Halpern et al., Nature, 2017; Inverso et al., Developmental Cell, 2021), and represents an advance over randomly distributed or non-patterned vascularized constructs.

The establishment of a functional microvascular network within the artificial liver scaffold was further supported by endothelial marker expression (CD31; Fig. 7) and perfusion assays, in which 5 µm fluorescent microparticles flowed freely through the endothelial-lined channels (Fig. 8b and Supplementary Video 1). The demonstrated perfusability is particularly significant, as convective transport through a patent lumen is a prerequisite for sustaining cell viability beyond the diffusion limit of approximately 100-200 µm and for scaling engineered tissues toward clinically relevant dimensions (Rouwkema & Khademhosseini, Trends in Biotechnology, 2016). These features are critical for sustaining tissue viability by facilitating nutrient and oxygen transport while enabling vascular network expansion (Devillard & Marquette, Frontiers in Bioengineering and Biotechnology, 2021; Pill et al., Frontiers in Bioengineering and Biotechnology, 2018; Pries & Secomb, Physiology, 2014). Consistent with this facilitative role, the scaffolds containing HG-EC aggregates and endothelial cells proliferated markedly faster than the HepG2-only group, reaching an approximately two-fold higher rate over seven days of culture (Fig. 6e). Endothelial cells were observed to migrate into and spread within the hepatocyte aggregates (Fig. 7h, j), providing direct morphological evidence of close spatial coupling between the two cell populations. In the native liver, sinusoidal endothelial cells release angiocrine factors, including hepatocyte growth factor and Wnt2, that promote hepatocyte proliferation and support hepatic homeostasis, and this angiocrine effect depends critically on the close physical proximity between endothelial cells and hepatocytes (Ding et al., Nature, 2010). The spatial architecture generated in our scaffolds recapitulates this proximity, providing a plausible mechanistic basis for both the enhanced proliferation and the improved hepatic function observed in the vascularized constructs (Fig. 9). In addition to this endothelial coupling, the liver-derived dECM matrix likely contributed to the observed hepatic function, consistent with reports that liver dECM enhances hepatocyte function relative to non-tissue-specific hydrogels (Sasaki et al., Biomaterials, 2017).

Consistent with this architecture-function relationship, the vascularized HG-EC scaffold demonstrated markedly enhanced hepatic marker expression and metabolic activity compared with the HepG2-only scaffold: by day 7, a 1.7-fold difference in urea production (Fig. 9a). The pronounced increase in cytochrome P450 enzyme expression (2.7-fold increase in *CYP1A2* and 6.2-fold increase in *CYP3A4*; Fig. 9d, g) is particularly relevant to hepatic maturation, as CYP3A4 is the most abundantly expressed drug-metabolizing enzyme in the adult human liver and its expression increases markedly during hepatic development from low prenatal levels (Wilkening et al., Drug Metab Dispos, 2003; Zanger & Schwab, Pharmacol Ther, 2013). The upregulation of CK18, an intermediate filament protein of mature hepatocytes, similarly supports improved hepatocyte organization (Fig. 9c). By contrast, the concurrent increase in AFP (Fig. 9b) should be interpreted with caution: AFP is a fetal hepatocyte marker whose expression normally declines with hepatic maturation and is constitutively expressed by HepG2 cells (He et al., Advanced Healthcare Materials, 2022). Its increase in our system therefore more likely reflects enhanced overall hepatocyte proliferation and biosynthetic activity than a shift toward a mature phenotype, and this distinction should be considered when interpreting the functional state of the constructs. Collectively, these results underscore the importance of coordinated vascular and parenchymal organization in promoting liver-specific function, while indicating that the constructs attain enhanced functional activity rather than full hepatocyte maturation.

The *in vivo* implantation experiments further demonstrate the translational potential of the engineered artificial liver scaffolds. Throughout the 21-day observation period, the implanted mice showed no signs of swelling, fever, or infection, with complete skin healing by day 7 and hair regrowth by day 21 (Fig. 2), indicating that both the surgical procedure and the scaffold itself were well tolerated. The explanted scaffolds also remained proliferative and viable over this entire 21-day period (Fig. 10a), indicating that the constructs retained functional cellular activity rather than merely persisting as an inert graft following transplantation. The early increase in inflammatory cytokines, which peaked on day 1 post-implantation in the scaffold-implantation group, likely reflects an acute immune response to the implanted scaffolds that resolved over the observation period, whereas the delayed cytokine peaks in the partial-hepatectomy group (day 14 for *IL-1*β and day 7 for *IL-16*) suggest a more prolonged tissue-injury response (de Jong et al., J Hepatol, 2001; Murtha-Lemekhova et al., Sci Rep, 2021; Söhling et al., Materials (Basel), 2022) (Fig. 10c, d). The comparatively early and transient inflammatory profile of the scaffold-implantation group is favorable, as a rapidly resolving acute response is generally more conducive to constructive tissue integration than a sustained inflammatory state. Importantly, the elevated expression of *TGF-*α in the scaffold-implantation group indicates active tissue remodeling and scaffold integration, processes that are critical for successful scaffold incorporation (Roshani et al., Cancer Letters, 2014; Sandgren et al., Mol Cell Biol, 1993) (Fig. 10b) and for maintaining normal hepatic function (Fig. 10e, f).

Histological analyses further confirmed the biocompatibility of the implanted scaffolds and their progressive integration with the host liver tissue. Evidence of vascular maturation was observed, as indicated by the progressive reduction in the diameter of the smallest blood-containing lumen, from 26.1 µm on day 7 to 7.9 µm by day 21, together with the presence of red blood cells within the vascular structures (Fig. 11). A comparable remodeling process, in which implanted engineered microvessels progressively shift toward smaller, more uniform diameters and become perfused with host-derived red blood cells, has been reported previously and interpreted as anastomosis with the host circulation (Cheng et al., Blood, 2011; Kang et al., Blood, 2011). While our histological data are consistent with the emergence of functional connectivity between the scaffold vasculature and the host circulation, two aspects remain to be confirmed. First, although the anucleate, biconcave morphology of the luminal contents is consistent with red blood cells, definitive identification would require erythroid-specific immunostaining, such as Ter-119. Second, the present study did not directly trace the origin of the luminal endothelium; the observed maturation may therefore result from remodeling of the implanted endothelial cells, ingrowth of host vessels, or a combination of both. Addressing these points will require erythroid-specific staining and lineage-tracing approaches in future work. Furthermore, Masson’s trichrome staining revealed a marked increase in collagen deposition within the grafted tissue (Fig. 12), reflecting dynamic extracellular matrix remodeling that likely contributes to stabilizing the scaffold–host interface.

Despite these promising findings, several limitations remain. First, the use of HepG2 cells, a hepatocellular carcinoma line, does not fully recapitulate the functional complexity of primary human hepatocytes, and longer-term *in vivo* studies are required to assess the durability and metabolic activity of the implanted scaffolds. Second, the EA.hy926 endothelial cell line, while experimentally tractable, does not fully reproduce the phenotype of primary endothelial cells, and in particular lacks the specialized features of liver sinusoidal endothelial cells that mediate angiocrine support within the lobule. Third, the current constructs do not incorporate other essential hepatic cell types, such as Kupffer cells, hepatic stellate cells, and cholangiocytes, that contribute to immune regulation, matrix homeostasis, and biliary function in the native liver. Fourth, while the engineered construct reconstitutes the basic microvascular–parenchymal unit of the hepatic lobule, it does not yet reproduce a complete lobule: the central-vein-scale draining vessel and the periportal-to-pericentral metabolic zonation were not recapitulated, and the vasculature comprises a relatively simple branched geometry.

Future work will therefore focus on incorporating primary or stem cell-derived hepatocytes and endothelial cells, integrating additional non-parenchymal cell types, and refining vascular-parenchymal interactions to further enhance functional outcomes. Toward larger, implantable-scale liver tissues, the present USW-patterned unit could potentially be integrated with our previously developed platforms for large-diameter vessel fabrication using an inverse-gravity nozzle (Duong et al., Biofabrication, 2023) and for multifascicle bundling of tissue units using a multi-barrel nozzle (Duong et al., Biomaterials, 2025); combining these approaches may enable a hierarchical construct in which a central-vein-scale supplying vessel connects to bundled, USW-patterned lobule-like units, although achieving uniform perfusion and functional integration across scales remains an important challenge. Given the demonstrated perfusability and hepatic functionality, the platform may also serve as a physiologically relevant model for drug metabolism and hepatotoxicity testing, in addition to its longer-term potential for implantable liver tissue engineering.

## 5. Conclusions

In this study, we developed a biofabrication approach for constructing artificial liver scaffolds that recapitulate the fundamental structural unit of liver tissue. Using two different frequency USW patterning, endothelial cells were arranged into branched vascular structures, which were then surrounded by HG-EC aggregates. The co-culture of endothelial cells and hepatocytes promoted the formation of a functional vascular network within the scaffold, which is essential for sustaining an adequate supply of oxygen and nutrients while facilitating the removal of metabolic waste. Moreover, the resulting vascular network supported hepatocyte proliferation and survival, thereby enhancing hepatocyte-specific function. The vascularized scaffolds exhibited markedly increased urea production and cytochrome P450 enzyme expression compared with hepatocyte-only cultures, indicating improved metabolic maturation. Following implantation into the mouse liver, the scaffolds showed favorable biocompatibility, progressive host integration, and evidence of vascular maturation, supporting their potential for *in vivo* application.

Collectively, these findings demonstrate that USW patterning provides a promising route toward the generation of fully vascularized artificial livers. This approach may have important implications for regenerative medicine and liver tissue engineering, potentially contributing to the development of functional hepatic grafts and helping to address the critical shortage of donor organs for patients with end-stage liver disease.

